# A Rapid, Cost-Effective Tailed Amplicon Method for Sequencing SARS-CoV-2

**DOI:** 10.1101/2020.05.11.088724

**Authors:** Daryl M. Gohl, John Garbe, Patrick Grady, Jerry Daniel, Ray H. B. Watson, Benjamin Auch, Andrew Nelson, Sophia Yohe, Kenneth B. Beckman

## Abstract

The global COVID-19 pandemic has led to an urgent need for scalable methods for clinical diagnostics and viral tracking. Next generation sequencing technologies have enabled large-scale genomic surveillance of SARS-CoV-2 as thousands of isolates are being sequenced around the world and deposited in public data repositories. A number of methods using both short- and long-read technologies are currently being applied for SARS-CoV-2 sequencing, including amplicon approaches, metagenomic methods, and sequence capture or enrichment methods. Given the small genome size, the ability to sequence SARS-CoV-2 at scale is limited by the cost and labor associated with making sequencing libraries. Here we describe a low-cost, streamlined, all amplicon-based method for sequencing SARS-CoV-2, which bypasses costly and time-consuming library preparation steps. We benchmark this tailed amplicon method against both the ARTIC amplicon protocol and sequence capture approaches and show that an optimized tailed amplicon approach achieves comparable amplicon balance, coverage metrics, and variant calls to the ARTIC v3 approach and represents a cost-effective and highly scalable method for SARS-CoV-2 sequencing.

## Introduction

The global COVID-19 pandemic has necessitated a massive public health response which has included implementation of society-wide distancing measures to limit viral transmission, the rapid development of qRT-PCR, antigen, and antibody diagnostic tests, as well as a world-wide research effort of unprecedented scope and speed. Next generation sequencing technologies (NGS) have recently enabled large-scale genomic surveillance of infectious diseases. Sequencing-based genomic surveillance has been applied to both endemic disease, such as seasonal influenza (1), and to emerging disease outbreaks such as Zika and Ebola (2–4).

As of May 2020, over 16,000 SARS-CoV-2 sequences have been deposited in public repositories such as NCBI and GISAID (5, 6). Several large-scale consortia in the UK (COG-UK: COVID-19 Genomics UK), Canada (CanCOGeN: Canadian COVID Genomics Network), and the United States (CDC SPHERES: SARS-CoV-2 Sequencing for Public Health Emergency Response, Epidemiology, and Surveillance) have begun coordinated efforts to sequence large numbers of SARS-CoV-2 genomes. Such genomic surveillance has already enabled insights into the origin and spread of SARS-CoV-2 (7, 8), including the sequencing efforts by the Seattle flu study which provided early evidence of extensive undetected community transmission of SARS-CoV-2 in the Seattle area (9).

A number of different approaches have been used to sequence SARS-CoV-2. Metagenomic (RNA) sequencing can be used to sequence and assemble SARS-CoV-2 (10). This approach has the disadvantage that samples must typically be sequenced very deeply in order to obtain sufficient coverage of the viral genome, and thus the cost of this approach is high relative to more targeted methods. Sequence capture methods (Figure 1A) can be used to enrich for viral sequences in order to lower sequencing costs and are being employed to sequence SARS-CoV-2 (11). Finally, amplicon approaches (Figure 1B), in which cDNA is made from SARS-CoV-2 positive samples and amplified using primers that generate tiled PCR products are being used to sequence SARS-CoV-2 (3). Since primers cannot capture the very ends of the viral genome, amplicon approaches have the drawback of slightly less complete genome coverage, and mutations in primer binding sites have the potential to disrupt the amplification of the associated amplicon. However, the relatively low-cost of amplicon methods make them a good choice for population-scale viral surveillance and such approaches have recently been used successfully to monitor the spread of viruses such as Zika and Ebola (2–4).

**Figure 1.**
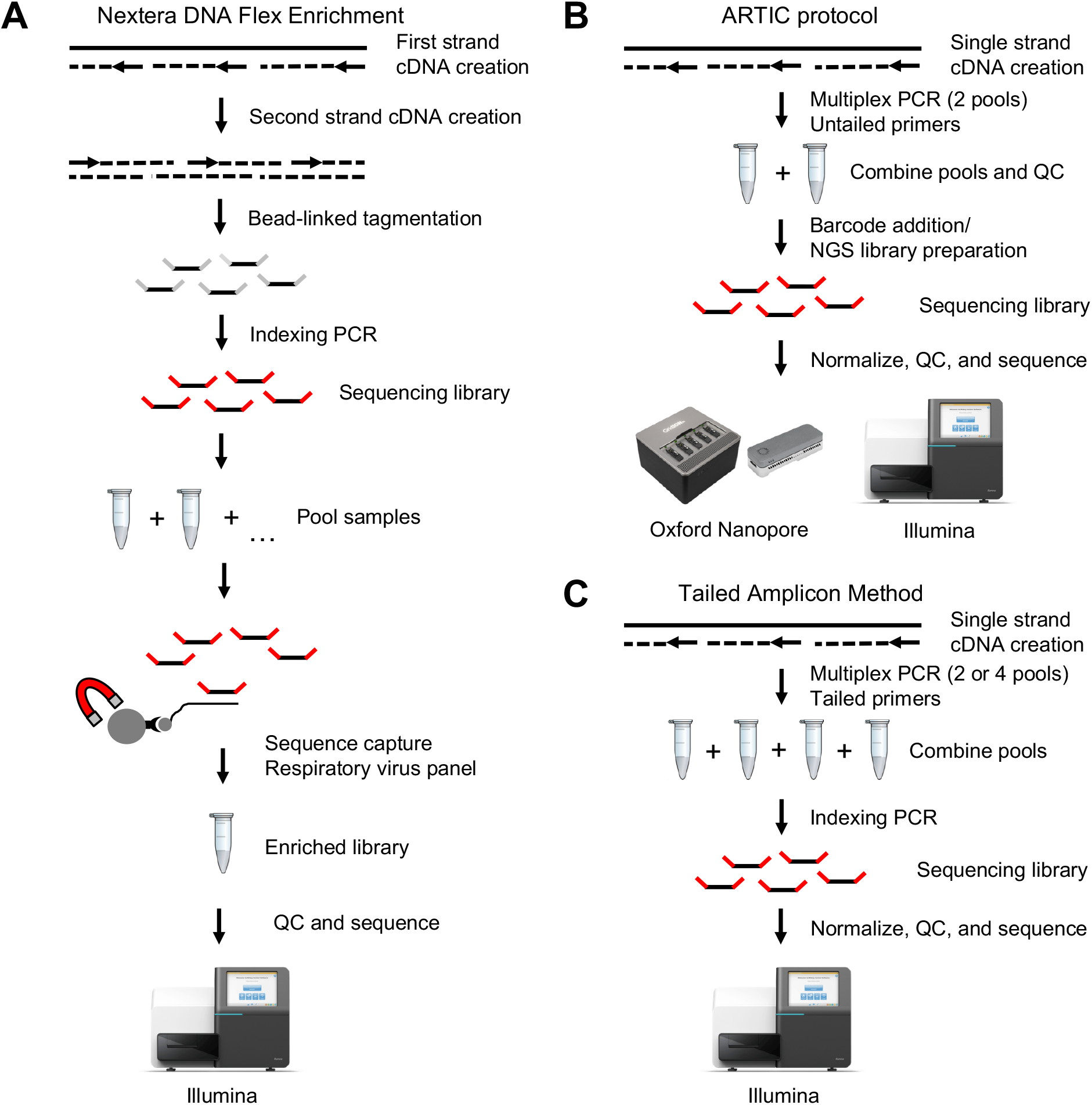
Methods for SARS-CoV-2 genome sequencing compared in this study. A) In Illumina’s Nextera DNA Flex Enrichment protocol cDNA is tagmented and made into barcoded sequencing libraries, which are then enriched using sequence capture with a respiratory virus panel containing probes against SARS-CoV-2. B) In the ARTIC protocol, first strand cDNA is enriched by amplifying with two pools of primers to generate amplicons tiling the SARS-CoV-2 genome. These amplicons are then subjected to either Illumina or Oxford Nanopore library preparation, using methods that either directly add adapters to the ends of the amplicons or fragment them to enable sequencing on a wider variety of Illumina instruments. C) The tailed amplicon approach, developed here, enriches first strand cDNA using ARTIC v3 primers containing adapter tails. This allows functional sequencing libraries to be created through a second indexing PCR reaction that adds sample-specific barcodes and flow cell adapters.

The ARTIC network (https://artic.network/) has established a method for preparing amplicon pools in order to sequence SARS-CoV-2 (Figure 1B). The ARTIC primer pools have gone through multiple iterations to improve evenness of coverage (12). Several variants of the ARTIC protocol exist in which the pooled SARS-CoV-2 amplicons from a sample are taken through a NGS library preparation protocol (using either ligation or tagmentation-based approaches) in which sample-specific barcodes are added, and are then sequenced using either short-read (Illumina) or long-read (Oxford Nanopore, PacBio) technologies. The library preparation step currently represents a bottleneck in sequencing SARS-CoV-2 amplicons, in terms of both cost and labor.

Here we describe an all-amplicon method for producing SARS-CoV-2 sequencing libraries which simplifies the process and lowers the per sample cost for sequencing SARS-CoV-2 genomes (Figure 1C). This approach incorporates adapter tails in the ARTIC v3 primer designs, allowing sequencing libraries to be produced in a two-step PCR process, bypassing costly and labor-intensive ligation or tagmentation-based library preparation steps. By re-optimizing the pooling strategy for the tailed primers, we demonstrate that this tailed amplicon approach can achieve similar coverage to the untailed ARTIC v3 primers at equivalent sequencing depths. We benchmark this approach against both the standard ARTIC v3 protocol and a sequence capture approach using clinical samples spanning a range of viral loads. The approach we describe is similar to a tailed-amplicon method that we have used to process more than 100,000 microbiome samples in recent years in the University of Minnesota Genomics Center (13), and thus represents a highly scalable method for sequencing large numbers of SARS-CoV-2 genomes in a rapid and cost-effective manner.

## Results

We designed a series of experiments in order to test a streamlined tailed amplicon method and to compare amplicon and sequence capture based methods for SARS-CoV-2 sequencing (Figure 1). We sequenced these samples using Illumina’s Nextera DNA Flex Enrichment protocol using a respiratory virus oligo panel containing probes for SARS-CoV-2, the ARTIC v3 tiled primers, and a novel tailed amplicon method designed to reduce cost and streamline the preparation of SARS-CoV-2 sequencing libraries.

We first evaluated the different SARS-CoV-2 sequencing workflows in their performance with a previously sequenced SARS-CoV-2 isolate strain from Washington state (2019-nCoV/USA-WA1/2020) provided by BEI Resources (14). As expected, since the amplicon approaches are unable to cover sequences at the ends of the SARS-CoV-2 genome, the DNA Flex Enrichment sequence capture method produced the highest genome coverage. At a subsampled read depth of 100,000 reads, the Nextera DNA Flex Enrichment method achieved 99.96% coverage at a minimum of 10x and 99.69% coverage at a minimum of 100x (Figure 2A-B). The ARTIC v3 method prepared with TruSeq library preparation achieved 99.60% coverage at a minimum of 10x and 97.31% coverage at a minimum of 100x (Figure 2A-B).

**Figure 2.**
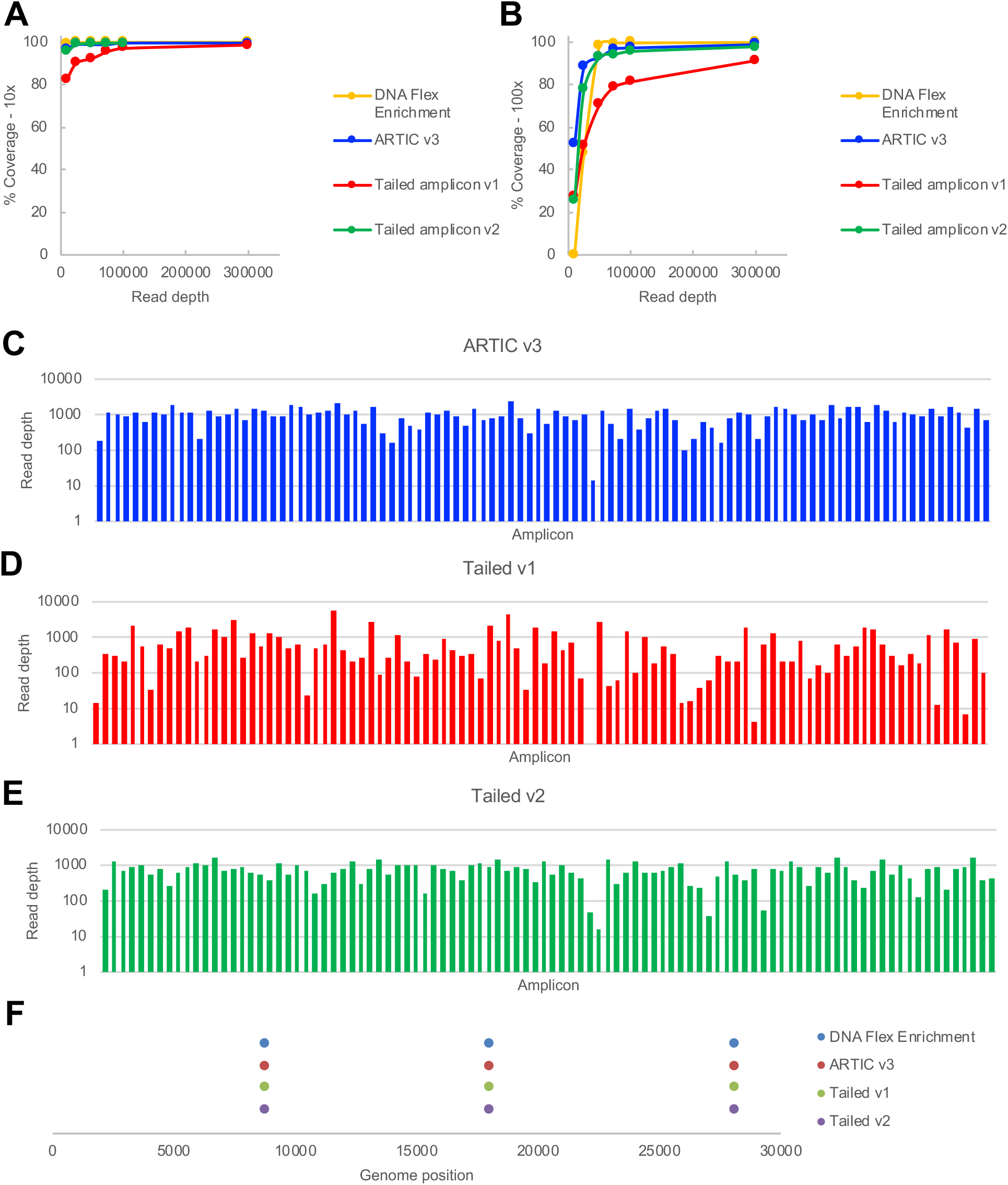
Comparison of sequence capture, ARTIC v3 amplicon, and tailed amplicon workflows on SARS-CoV-2 isolate. A) Percentage of the BEI WA isolate genome coverage at 10x at different subsampled read depths when sequenced with the indicated approach. B) Percent of the BEI WA isolate genome coverage at 100x at different subsampled read depths when sequenced with the indicated approach. C) Observed read depth for each of the expected amplicons for the BEI WA isolate amplified with the ARTIC v3 protocol at a subsampled read depth of 100,000 raw reads. D) Observed read depth for each of the expected amplicons for the BEI WA isolate amplified with the tailed amplicon v1 (2 pool amplification) protocol at a subsampled read depth of 100,000 raw reads. E) Observed read depth for each of the expected amplicons for the BEI WA isolate amplified with the tailed amplicon v2 protocol (4 pool amplification) at a subsampled read depth of 100,000 raw reads. F) Positions of variants detected for the BEI WA isolate at a read depth of up to 1,000,000 raw reads (or the maximum read depth for the sample).

We tested a tailed amplicon method (tailed amplicon v1) in which the tailed version of the ARTIC v3 primers were pooled into two pools in a similar manner to the ARTIC v3 protocol. The BEI WA isolate strain was amplified for both 25 or 35 PCR cycles, using the same enzymes and PCR conditions used for the ARTIC v3 data set. The tailed amplicon v1 method produced lower coverage than the ARTIC v3 method, with 98.87% coverage at a minimum of 10x and 89.40% coverage at a minimum of 100x for the 25 PCR cycle sample and 97.09% coverage at a minimum of 10x and 81.31% coverage at a minimum of 100x for the 35 PCR cycle sample (Figure 2A-B). The poorer performance with respect to coverage metrics with the tailed amplicon v1 protocol was due to substantially worse balance between the different tiled amplicons than with the ARTIC v3 (untailed) primers (Figure 2C-D). The coefficient of variation (CV) of the ARTIC v3 sample was 0.49 and the CVs of the tailed amplicon v1 samples were 1.70 and 1.26 for the 25 and 35 PCR cycle samples, respectively.

The ARTIC v3 primers have been through multiple cycles of iteration to achieve relatively even amplicon balance and genome coverage (12). We reasoned that reducing the concentration of the primers that were over-represented in the initial round of sequencing may improve balance. While adjusting the primer concentration for over-represented amplicons did lower the CV of the tailed amplicon pool, amplicon balance was still substantially worse than with the untailed ARTIC v3 primers (data not shown).

We next tested whether splitting the tailed SARS-CoV-2 primers into 4 PCR reactions based on primer performance in the initial sequencing tests could improve balance with the tailed primer approach. The 4-pool amplification scheme (tailed amplicon v2) achieved coverage metrics close to the untailed ARTIC v3 approach at comparable read depths with 98.76% coverage at a minimum of 10x and 95.64% coverage at a minimum of 100x (Figure 2A-B). The improvement in genome coverage metrics with the tailed amplicon v2 approach was a function of improved amplicon balance (Figure 2E). The CV of the tailed amplicon v2 sample was 0.52 (comparable to the CV of 0.49 with the untailed ARTIC v3 approach). The same three variants were detected by all four methods tested (Figure 2F), consistent with prior comparisons of the USA-WA1/2020 and the Wuhan-Hu-1 reference strain.

Next, we assessed the performance of the different SARS-CoV-2 sequencing approaches on a set of deidentified patient samples. We selected 9 SARS-CoV-2 positive patient samples spanning a range of viral loads as assessed by a qRT-PCR using the CDC primers targeting the SARS-CoV-2 nucleocapsid gene (N1 and N2 targets, Supplemental Figure 1). In addition, we included two patient negative samples in these experiments. We carried out initial tests of the Nextera DNA Flex Enrichment protocol, the tailed amplicon v1 approach, and the ARTIC v3 approach using this sample set. For testing the tailed amplicon v2 approach, and comparing among all four methods, we used a subset of these patient samples with N1 and N2 Ct values ranging from ~20-35 (Figure 3A).

**Figure 3.**
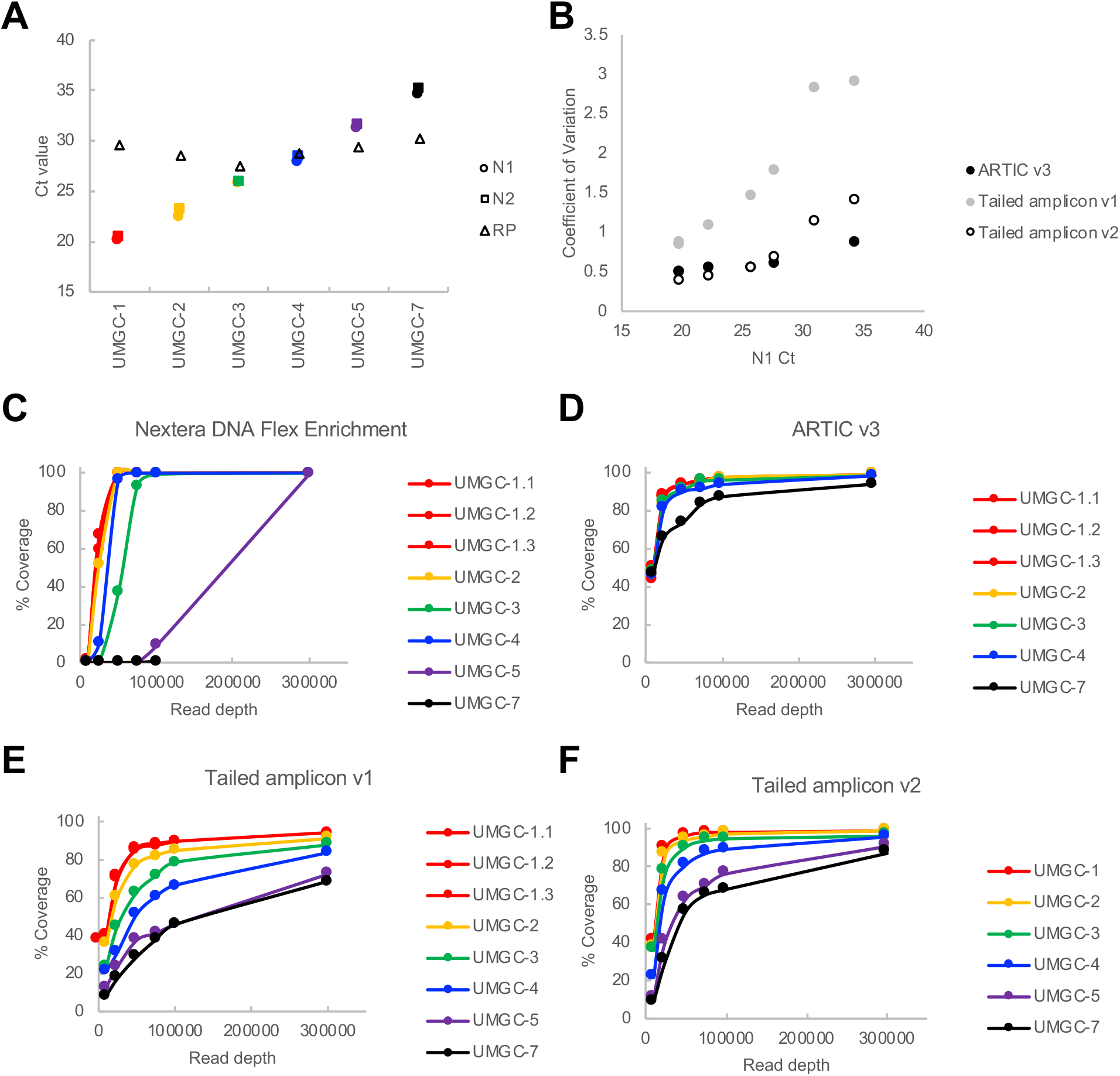
Comparison of sequence capture, ARTIC v3 amplicon, and tailed amplicon workflows on clinical specimens spanning a range of viral loads. A) Samples with N1 and N2 Ct values ranging from approximately 20-35 chosen for testing of SARS-CoV-2 sequencing workflows. B) Evenness of representation of amplicons for different workflows as a function of sample N1 Ct value. Percentage of genome coverage at 100x at different subsampled read depths for each sample when sequenced using the following approaches: D) Illumina Nextera DNA Enrichment; E) ARTIC v3 with TruSeq library preparation. F) Tailed amplicon v1 (2 pool amplification); G) Tailed amplicon v2 (4 pool amplification)

For the Illumina DNA Flex Enrichment protocol, SARS-CoV-2 genome coverage was more complete for samples with lower N1 and N2 Cts (ranging from ~20-30) at comparable read depths and coverage thresholds than with amplicon approaches, similar to the BEI WA isolate data (Figure 3C, Supplemental Figure S2-S3). However, for samples with N1 and N2 Ct values greater than approximately 30, the number of sequencing reads were substantially reduced and the proportion of reads mapping to the human genome were substantially increased (Supplemental Figure S4). The average coverage at a subsampled read depth of 100,000 raw reads was 99.86% (10x) and 67.94% (100x) for all six test samples. For samples with N1 and N2 Ct vales of less than 30, average coverage was 99.94% (10x) and 98.01% (100x) at a subsampled read depth of 100,000 raw reads.

For ARTIC v3 tests, based on the N1 and N2 target Ct values from clinical testing, we used either 25, 30, or 35 PCR cycles for the amplification reactions. Sufficient amplification to carry out TruSeq library prep was seen for samples with Cts of around 35 or less. Five patient samples with N1 and N2 Ct values ranging from ~20-35 and the BEI WA isolate sample were selected for TruSeq library prep and sequencing; one sample (N1 Ct = 20, N2 Ct = 20.4) was prepared in triplicate. Consistent with previous descriptions of the ARTIC v3 primers, the balance between the tiled amplicons across these samples was relatively even, with a mean CV of 0.61 among the five patient samples tested, and 0.55 for samples with a N1 and N2 Ct of less than 30 (Figure 3B, Supplemental Figure S5). For the ARTIC v3 protocol, the average coverage at a subsampled read depth of 100,000 raw reads was 98.87% (10x) and 94.21% (100x) for all five test samples. For samples with N1 and N2 Ct vales of less than 30, average coverage was 98.87% (10x) and 95.95% (100x) at a subsampled read depth of 100,000 raw reads (Figure 3D, Supplemental Figure S2-S3).

We performed initial tests of the tailed amplicon v1 protocol by amplifying the samples listed in Figure 3A for 25 or 35 PCR cycles using tailed versions of the ARTIC v3 primers split into two separate pools. As with the BEI WA isolate sample, the balance observed with the tailed amplicon v1 approach was worse than the ARTIC v3 protocol, with a mean CV of 1.81 among the six patient samples tested, and 1.28 for samples with a N1 and N2 Ct of less than 30 (Figure 3B, Supplemental Figure S6). This led to decreased coverage at a given read depth for the tailed amplicon v1 method relative to ARTIC v3 (Figure 3E, Supplemental Figure S2).

Upon splitting the tailed SARS-CoV-2 primers into 4 PCR reactions based on primer performance in the initial sequencing tests, the tailed amplicon v2 method had much improved amplicon balance. The mean CV of all six patient samples was 0.76 (compared to a CV of 0.61 with ARTIC v3) and 0.52 for samples with a N1 and N2 Ct of less than 30 (compared to 0.55 with the ARTIC v3 protocol; Figure 3B, Supplemental Figure S7). The tailed amplicon v2 protocol had an average coverage at a subsampled read depth of 100,000 raw reads of 98.60% (10x) and 87.17% (100x) for all six test samples. For samples with Ct vales of less than 30, average coverage was 98.81% (10x) and 94.72% (100x) at a subsampled read depth of 100,000 raw reads (Figure 3F, Supplemental Figure S2-S3).

The slightly lower coverage metrics at a given subsampled read depth for the tailed amplicon v2 method can likely be explained by primer dimer formation during the two-step amplification process, which is more pronounced for higher N1 and N2 Ct samples (Supplemental Figure S8). Despite observing negligible amounts of primer dimer products on the bioanalyzer trace, samples with N1 and N2 Ct values greater than 30 had as much as 50% primer dimer in the resulting sequencing reads. We have previously reported a substantial size bias on the MiSeq, which may help explain the preferential clustering and out-sized proportion of primer dimer reads present in the sequencing data for some samples (15). While this issue can be overcome by increased sequencing depth, future optimizations aimed at reducing primer dimer contamination such as more stringent size selection or sequencing on an instrument with less size bias, such as the NovaSeq (15) could reduce this effect.

Finally, we examined the variants detected in the patient samples for each of the SARS-CoV-2 sequencing methods. There was complete concordance in the variant calls for all samples with N1 and N2 Ct values below 30, but less agreement among variant calls between methods for the sample with N1 and N2 Ct values of approximately 35 (Figure 4).

**Figure 4.**
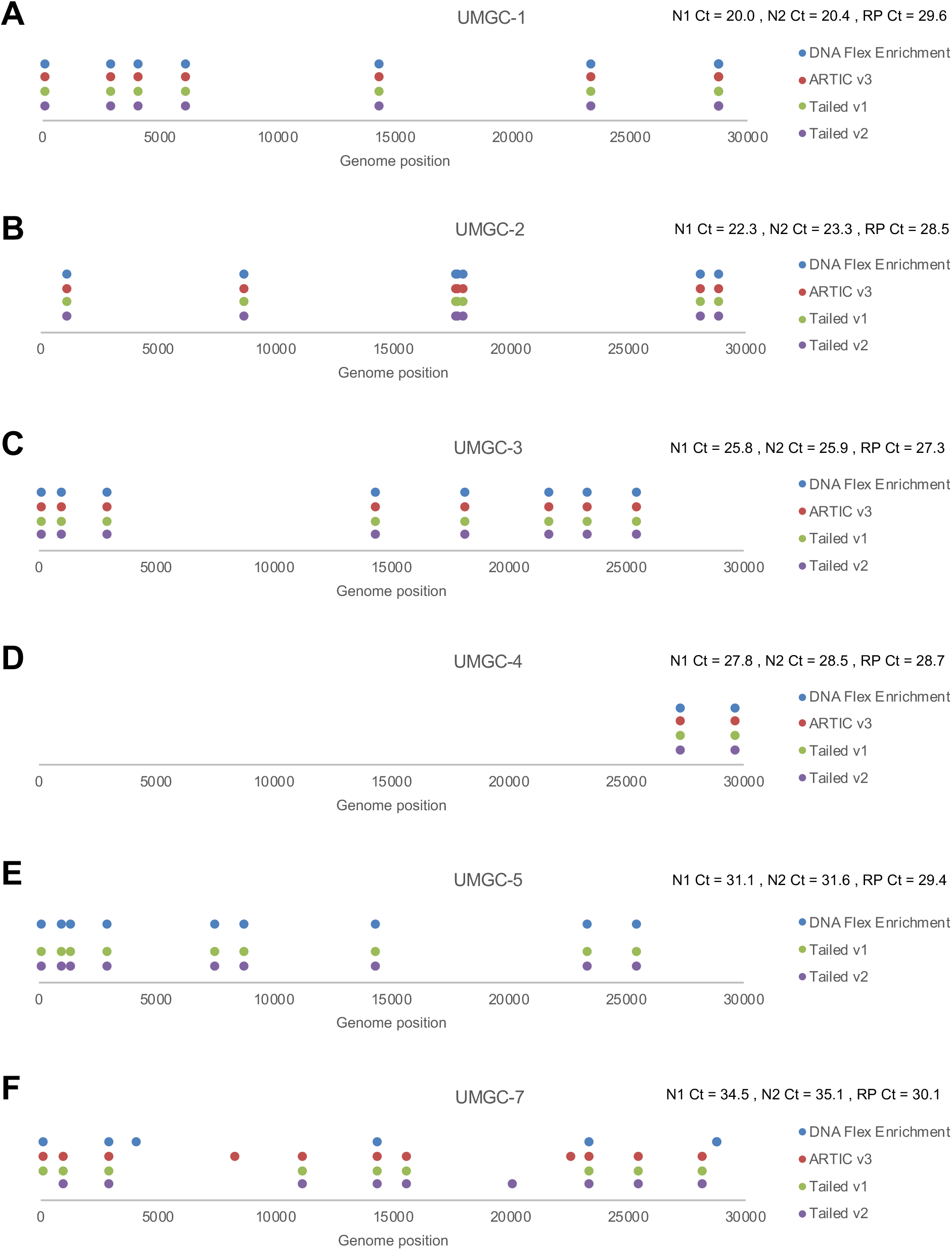
Variants detected using different sequencing workflows. A-F) Positions of variants detected for the indicated and sequencing protocol at a read depth of up to 1,000,000 raw reads (or the maximum read depth for the sample).

## Discussion

Here we compare sequence capture and amplicon-based methods for sequencing SARS-CoV-2 and describe a streamlined tailed amplicon method for cost-effective and highly scalable SARS-CoV-2 sequencing. In comparing the sequence capture and amplicon-based methods, there is a trade-off between the completeness of genome coverage and sensitivity (being able to analyze samples with higher N1 and N2 Ct values. Consistent with other recent analyses of SARS-CoV-2 amplicon sequencing approaches (16), we observed highly concordant results from samples with N1 and N2 Ct values of less than 30. For samples with Ct values between 30 and 35, coverage metrics tended to be less robust at a given read depth and samples with Ct values of greater than 35 did not perform well under any of the conditions tested. Based on validation experiments for the University of Minnesota qRT-PCR clinical COVID-19 diagnostic assay, we estimate that a Ct value of 30 corresponds to roughly 500 SARS-CoV-2 genome copies and a Ct value of 35 corresponds to roughly 15 SARS-CoV-2 genome copies in the 5 μL input used for cDNA creation (17).

We describe a modified workflow for SARS-CoV-2 sequencing which builds on the tiled amplicon approach developed by the ARTIC consortium and currently employed by many labs around the world. This tailed amplicon method uses a two-step PCR process similar to workflows previously described by us and others to generate microbiome or other amplicon sequencing data (13). Through an iterative testing process, we demonstrate that with the tailed amplicon v2 method, a four-pool amplification scheme produces data with comparable amplicon balance, coverage metrics, and variant calls to the ARTIC v3 approach. The tailed amplicon approach bypasses costly and labor-intensive library preparation steps and will allow for production of SARS-CoV-2 libraries at high scale (similar workflows are run on tens of thousands of samples per year in the University of Minnesota Genomics Center) at low cost (between $20-40 per sample depending on scale, including labor costs). We anticipate that this approach will aid in the genomic surveillance of SARS-CoV-2 as well as studies on viral diversity and evolution, and the influence of virus genetics on transmissibility, virulence, and clinical outcomes.

## Methods

### Samples

Extracted RNA from de-identified clinical biospecimens were obtained subsequent to COVID-19 testing at the University of Minnesota for use under the IRB approved protocol “Detection of COVID 19 by Molecular Methods” (STUDY00009560). Nine samples spanning a range of viral loads as assessed by the Ct values of the viral N1 and N2 targets by qRT-PCR were selected for these studies. In addition, two SARS-CoV-2 negative samples were selected to assess cross-contamination or other sequencing artifacts. The following reagent was deposited by the Centers for Disease Control and Prevention and obtained through BEI Resources, NIAID, NIH: Genomic RNA from SARS-Related Coronavirus 2, Isolate USA-WA1/2020, NR-52285.

### RNA extraction

RNA was extracted using one of three kits (Qiagen QIAamp Viral RNA Mini kit, Macherey-Nagel Nucelospin Virus Mini kit, and Biomérieux easyMag NucliSENS system) as described previously (17). All extraction methods used 100 μL of viral transport medium as input and eluted in 100 μL of appropriate elution buffer as indicated by manufacturer protocols. The integrity of the extracted RNA was analyzed using the Agilent high sensitivity RNA screentape assay on Agilent 2200 TapeStation following the manufacturer’s guidelines (Agilent, Santa Clara, CA).

### SARS-CoV-2 qRT-PCR

qRT-PCR reactions to identify SARS-CoV-2 samples were carried out using a modified version of the Centers for Disease Control and Prevention (CDC) SARS-CoV-2 qRT-PCR assay, as previously described (17). Briefly, three separate 10 μL RT-qPCR reactions were set up in a 384-well Barcoded plate (Thermo Fisher Scientific, Waltham, MA) for either the N1, N2, or RP primers and probes. 2.5 μL extracted RNA was added to 7.5 μL qPCR master mix comprised of the following components: 1.55 μL nuclease-free water, 5 μL GoTaq^®^ Probe qPCR Master Mix with dUTP (2X) (Promega, Madison, WI), 0.2 μL GoScript™ RT Mix for 1-Step RT-qPCR (Promega, Madison, WI), 0.75 μL primer/probe sets for either N1, N2, or RP (IDT, Coralville, IA). Reactions were run on a QuantStudio QS5 (Thermo Fisher Scientific, Waltham, MA) using the following cycling conditions: one cycle of 45°C for 15 minutes, followed by one cycle of 95°C for 2 minutes, followed by 45 cycles of 95°C for 15 seconds and 60°C for 1 minute. A minimum of two no template controls (NTCs) were included on all runs. A ΔRn threshold of 0.5 was selected and set uniformly for all runs. Ct values were exported and analyzed in Microsoft Excel.

### ARTIC v3 amplicon library generation and sequencing

The following reaction was set up to create cDNA using the ARTIC v3 protocol: 5 μL template RNA, 11 μL nuclease-free water, 4 μL SuperScript IV VILO master mix (Thermo Fisher Scientific, Waltham, MA). cDNA synthesis reactions were incubated at: 25°C for 10 minutes, followed by 50°C for 10 minutes and 85°C for 5 minutes. cDNA was amplified using each of the two ARTIC v3 primer pools which tile the SARS-CoV-2 genome. The following recipe was used to set up the PCR reactions: 2.5 μL template cDNA, 14.75 μL nuclease-free water, 5 μl 5x Q5 reaction buffer (New England Biolabs, Ipswich, MA), 0.5 μL 10 mM dNTPs (Kapa Biosystems, Woburn, MA), 0.25 μL Q5 Polymerase (New England Biolabs, Ipswich, MA), 2 μL primer pool 1 or 2 (10 μM). Cycling conditions were: 98°C for 30 seconds, followed by 25 or 35 cycles of 98°C for 15 seconds and 65°C for 5 minutes. Pools 1 and 2 were then combined, cleaned up with 1:1 AMPureXP beads (Beckman Coulter, Brea, CA)., and quantified by Qubit Fluorometer and Broad Range DNA assay (Thermo Fisher Scientific, Waltham, MA) and TapeStation capillary electrophoresis (Agilent, Santa Clara, CA).

Eight samples with >1ng/μL concentration of target amplicons were selected for downstream library preparation. Library preparation was performed following the standard Illumina TruSeq Nano DNA protocol for 350 base pair libraries (Illumina, San Diego, CA). A total of 100ng of amplicons from the ARTIC protocol were used as the input for library preparation. Input material was not sheared, as the amplicons were already the desired fragment length.

### Nextera DNA Flex Enrichment with respiratory virus panel

A modified non-directional NEBNext Ultra II First and Second Strand (#E7771 and #E6111, New England Biolabs, Ipswich, MA) protocol was used to generate long fragments of double-stranded cDNA as input material for the Nextera DNA Flex Enrichment with respiratory virus panel. The following reaction was set up for non-fragmented priming of RNA: 5 μL template RNA and 1 μL NEBNext Random Primers were combined and incubated at 65°C for 5 minutes. Non-directional first strand cDNA synthesis was performed by combining 6 μl of primed template RNA, 4 μL NEBNext First Strand Synthesis Buffer, 2 μL NEBNext First Stand Synthesis Enzyme Mix, and 8 μL nuclease-free water. The first strand synthesis reaction was incubated at 25°C for 10minutes, 42°C for 50 minutes, 70°C for 15 minutes. Second strand cDNA synthesis was performed by combining 20 μl first strand synthesis product, 8 μL of NEBNext Second Strand Synthesis Reaction Buffer with dUTP mix (10X), 4 μL NEBNext Second Strand Synthesis Enzyme Mix, and 48 μL nuclease-free water. The second strand synthesis reaction was incubated at 16°C for 60 minutes. Double-stranded cDNA was purified and concentrated with 1.8X AMPureXP beads (Beckman Coulter, Brea, CA) before eluting into 30 μL of 0.1X TE Buffer. Double-stranded cDNA size was determined using Bioanalyzer high sensitivity DNA assay (Agilent, Santa Clara, CA) and quantified with Qubit Fluorometer and High Sensitivity DNA assay (Thermo Fisher Scientific, Waltham, MA). cDNA was used to generate libraries using the Nextera DNA Flex Enrichment protocol (Illumina, San Diego, CA, catalog number 20025524) with the respiratory virus oligo panel including SARS-CoV-2 probes (Illumina, San Diego, CA, catalog number 20042472) according to manufacturer’s instructions.

### Two-pool tailed amplicon library generation and sequencing

To generate cDNA upstream of SARS-CoV-2 genome amplification, the following reaction was set up: 5 μL template RNA, 11 μL nuclease-free water, 4 μL SuperScript IV VILO master mix (Thermo Fisher Scientific, Waltham, MA). cDNA synthesis reactions were incubated at: 25°C for 10 minutes, followed by 50°C for 10 minutes and 85°C for 5 minutes. The SARS-CoV-2 genome was amplified using a two-step PCR protocol. The primary amplification was carried out in a manner similar to the ARTIC v3 method described above, using two primer pools which tile the SARS-CoV-2 genome. The following recipe was used to set up the PCR reactions: 2.5 μL template cDNA, 14.75 μL nuclease-free water, 5 μL 5x Q5 reaction buffer (New England Biolabs, Ipswich, MA), 0.5 μL 10 mM dNTPs (Kapa Biosystems, Woburn, MA), 0.25 μL Q5 Polymerase (New England Biolabs, Ipswich, MA), 2 μL primer pool 1 or 2 (10 μM) for the tailed v1 protocol. Cycling conditions were: 98°C for 30 seconds, followed by 25 or 35 cycles of 98°C for 15 seconds and 65°C for 5 minutes. The primers for the primary amplification contained both SARS-CoV-2 targeting sequences (derived from the ARTIC v3 designs), as well as adapter tails for adding indices and Illumina flow cell adapters in a secondary amplification. These amplification primers had the following structure (see Supplementary Information for primer sequences):

Left primers: TCGTCGGCAGCGTCAGATGTGTATAAGAGACAG**<SARS-CoV-2 LEFT primer>**
Right primers: GTCTCGTGGGCTCGGAGATGTGTATAAGAGACAG**<SARS-CoV-2 RIGHT primer>**

The PCR products from pool 1 and pool 2 for each sample were combined and then diluted 1:100 in sterile, nuclease-free water, and a second PCR reaction was set up to add the Illumina flow cell adapters and indices. The secondary amplification was done using the following recipe: 5 μL template DNA (1:100 dilution of the first PCR reaction), 0.7 μL nuclease-free water, 2 μL 5x Q5 reaction buffer (New England Biolabs, Ipswich, MA), 0.2 μL 10 mM dNTPs (Kapa Biosystems, Woburn, MA, 0.1 μL Q5 Polymerase (New England Biolabs, Ipswich, MA), 0.5 μL forward primer (10 μM), 0.5 μL reverse primer (10 μM). Cycling conditions were: 98°C for 30 seconds, followed by 10 cycles of 98°C for 20 seconds, 55°C for 15 seconds, 72°C for 1 minute, followed by a final extension at 72°C for 5 minutes. The following indexing primers were used (X indicates the positions of the 10 bp unique dual indices):

Forward indexing primer: AATGATACGGCGACCACCGAGATCTACACXXXXXXXXXXTCGTCGGCAGCGTC
Reverse indexing primer: CAAGCAGAAGACGGCATACGAGATXXXXXXXXXXGTCTCGTGGGCTCGG

### Four-pool tailed amplicon v2 library generation and sequencing

Samples were processed as described above for the two-pool tailed amplicon sequencing workflow, with the exception that in the first round of PCR, four separate reactions were set up using primer pools 1.1, 1.2, 2.1, and 2.2 (see Supplementary Information for primer sequences). The four PCR reactions were combined in a 1:1:1:1 ratio after an initial PCR amplification of 35 cycles and a 1:100 dilution of the combined PCRs for each sample was indexed according to the process described above.

### Normalization and pooling of tailed amplicon sequencing libraries

10 μL of PCR product for each sample was normalized using a SequalPrep 96-well Normalization Plate Kit (Thermo Fisher Scientific, Waltham, MA). Samples were eluted in 20 μL of elution buffer and 10 μL of each sample was pooled and concentrated to 20 μL using 0.7x AMPureXP beads (Beckman Coulter, Brea, CA). The final pooled sample was quantified using a Qubit Fluorometer and High Sensitivity DNA assay (Thermo Fisher Scientific, Waltham, MA). To confirm the expected library size of approximately 550 bp, pooled libraries were run on either an Agilent Bioanalyzer or TapeStation (Agilent, Santa Clara, CA).

### Sequencing

The sample pools were diluted to 2 nM based on the Qubit measurements and Agilent sizing information, and 10 μL of the 2 nM pool was denatured with 10 μL of 0.2 N NaOH. Amplicon libraries (ARTIC v3, Tailed v1, Tailed v2) were diluted to 8 pM in Illumina’s HT1 buffer, spiked with 5% PhiX, and sequenced using a MiSeq 600 cycle v3 kit (Illumina, San Diego, CA). The Nextera DNA Flex Enrichment library was diluted to 10 pM in Illumina’s HT1 buffer, spiked with 1% PhiX, and sequenced using a and a MiSeq 300 cycle v2 kit (Illumina, San Diego, CA).

### Analysis

The analysis method for amplicon libraries is as follows: Sample quality was assessed with FastQC (18). Read-pairs were stitched together using PEAR (19). Human host DNA was filtered by aligning the stitched reads to the human genome (GRCh38). Reads that did not align to the host genome were aligned to the reference Wuhan-Hu-1 (5) SARS-CoV-2 genome (MN908947.3) using BWA (20). Amplicon read depths were determined by counting the number of aligned reads covering the base at the center of each amplicon region. The iVar software package was used to trim primer sequences from the aligned reads, and iVar and Samtools mpileup were used to call variants and generate consensus sequences (3). Variants located outside of the region targeted by the amplicon panel were filtered out (reference genome positions 1-54 and 29836-29903), and consensus sequences bases corresponding to those regions were trimmed. The Nextera DNA Flex Enrichment libraries were analyzed using the same process, except the iVar primer trimming step was omitted, and no filtering of variants or trimming of consensus sequence was performed.

## Data Availability

Sequencing data for this project is available through the NCBI Sequence Read Archive BioProject PRJNA631042. Genome sequences of the strains sequenced in this study are available in GenBank BioProject PRJNA631042.

## Acknowledgements

We thank the staff of the University of Minnesota Genomics Center for helpful discussions and technical support. We thank Brandon Vanderbush for conducting QC on the SARS-CoV-2 samples and sequencing libraries. This work was carried out in part using computing resources at the University of Minnesota Supercomputing Institute. We thank Sean Wang and Matt Plumb from the Minnesota Department of Heath for helpful discussions and for sharing ARTIC v3 primers.

## Author contributions

D.M.G. conceived and designed the experiments, conducted experiments, analyzed data, and wrote the manuscript; K.B.B. conceived and designed the experiments and helped write the manuscript; J.G. analyzed data and helped write the manuscript; P.G., J.D., R.W., and B.A. conducted the experiments and helped write the manuscript; A.N. and S.Y. contributed experimental samples and helped write the manuscript.

## Supplemental Figure Legends

**Supplemental Figure S1.**
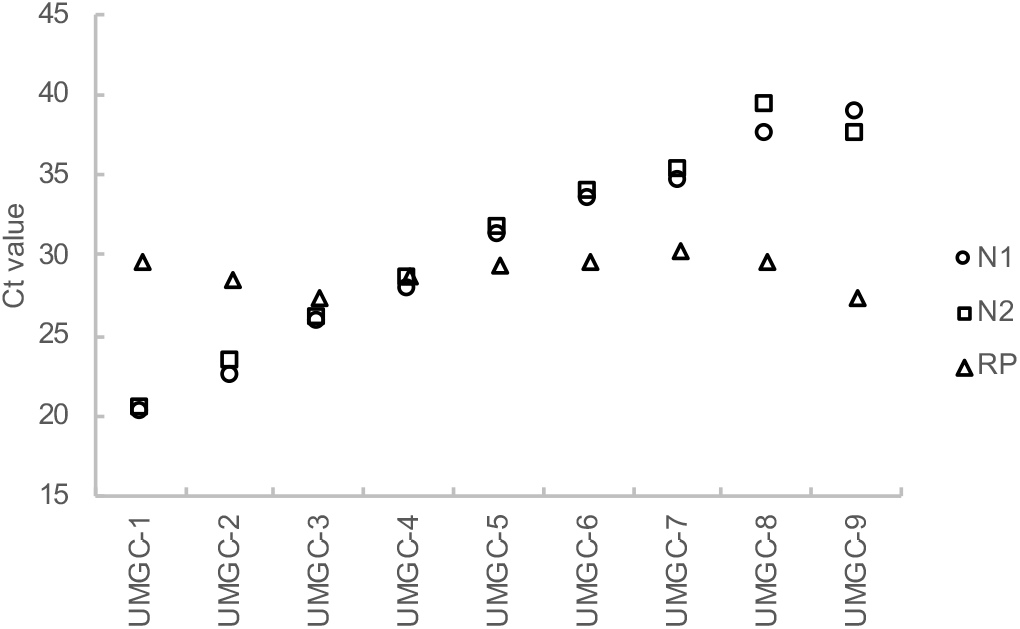
Samples for initial SARS-CoV-2 sequencing workflow tests. Samples with N1 and N2 Ct values ranging from approximately 20-35 chosen for testing of SARS-CoV-2 sequencing workflows.

**Supplemental Figure S2.**
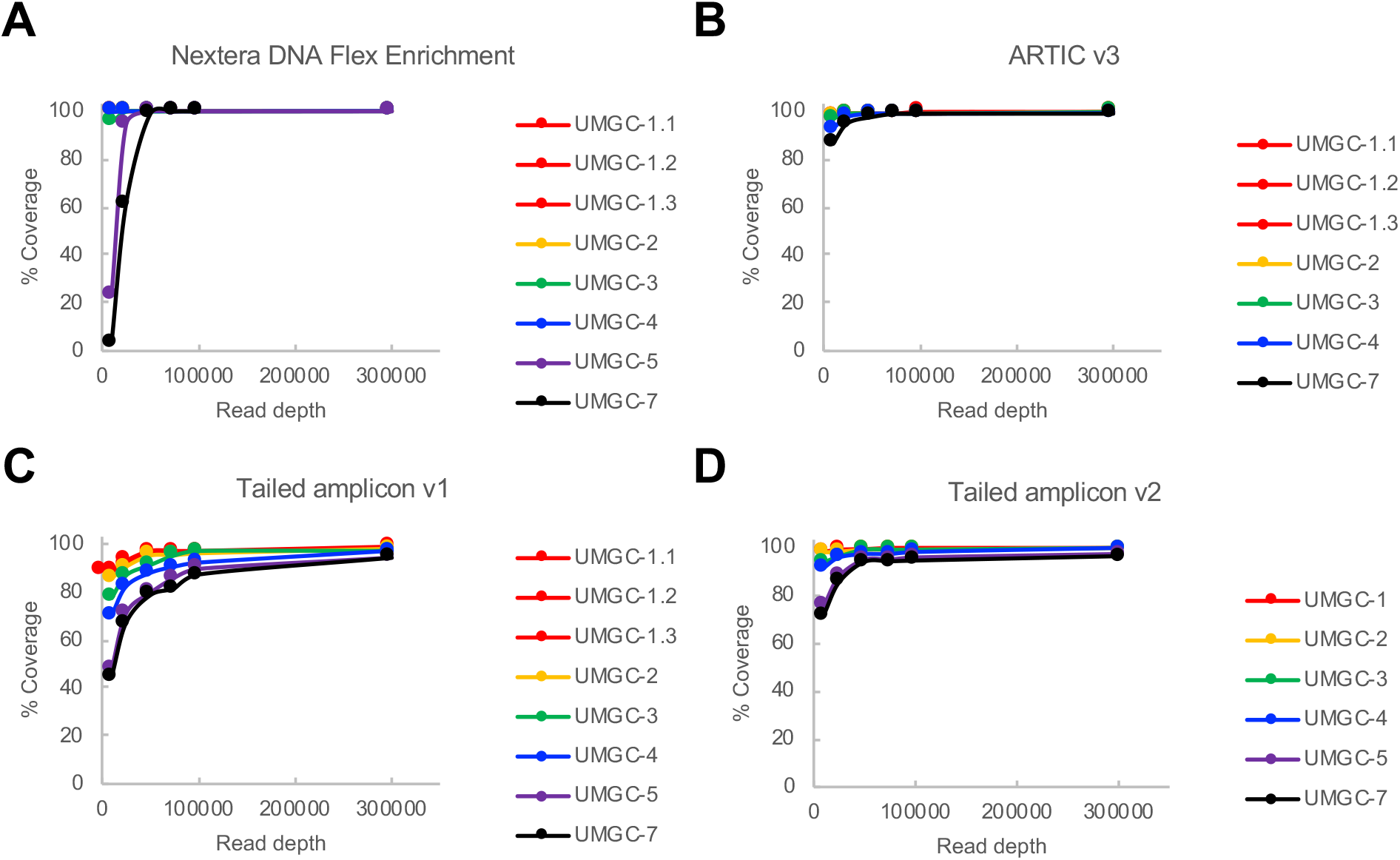
Coverage metrics by method for sequence capture, ARTIC v3 amplicon, and tailed amplicon workflows. Percentage of genome coverage at 10x at different subsampled read depths for each sample when sequenced using the following approaches: A) Illumina Nextera DNA Enrichment; B) ARTIC v3 with TruSeq library preparation. C) Tailed amplicon v1 (2 pool amplification); D) Tailed amplicon v2 (4 pool amplification).

**Supplemental Figure S3.**
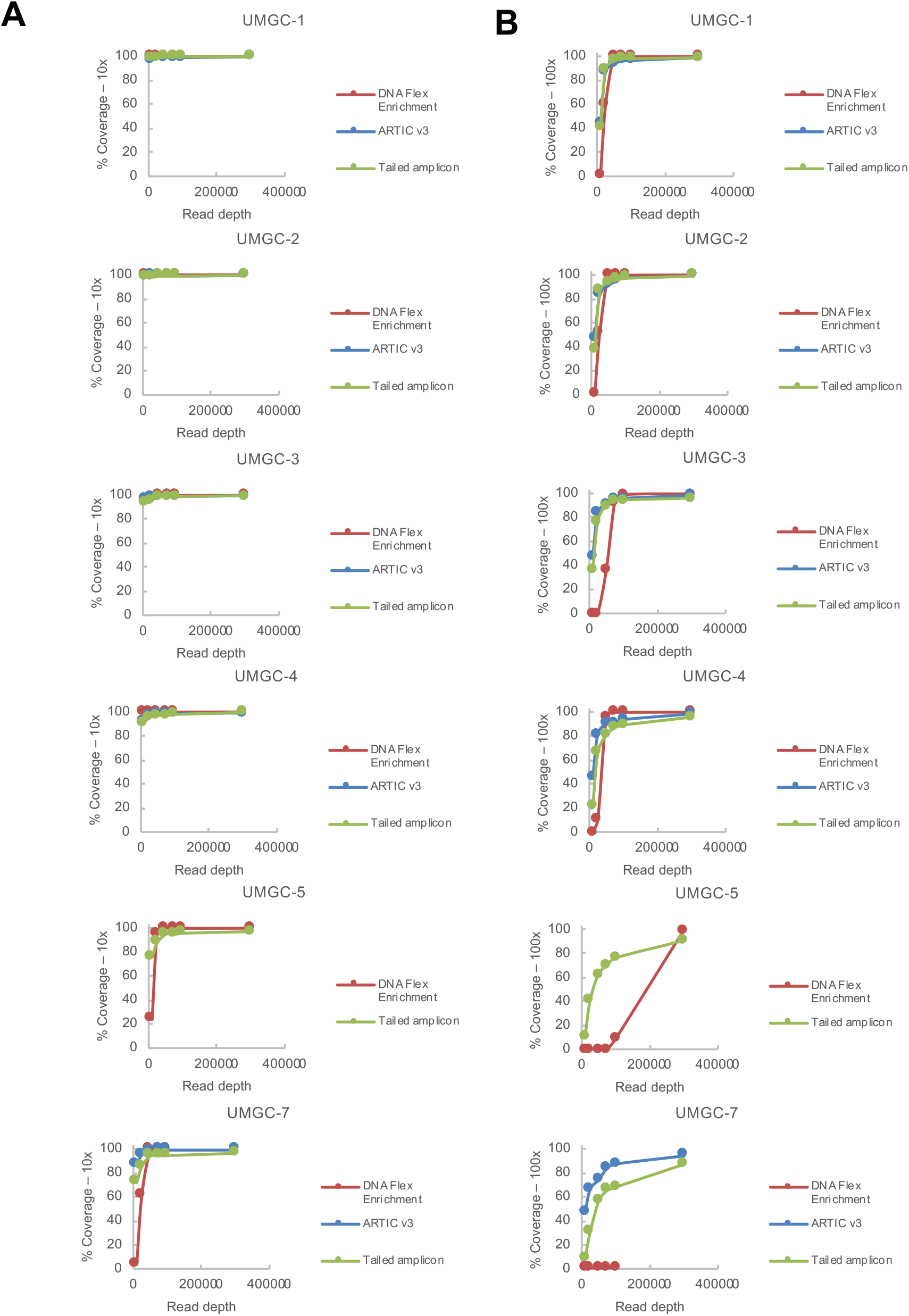
Coverage metrics by sample for sequence capture, ARTIC v3 amplicon, and tailed amplicon workflows. A) Percentage of genome coverage at 10x at different subsampled read depths for the indicated sample when sequenced using the indicated workflow. B) Percentage of genome coverage at 100x at different subsampled read depths for the indicated sample when sequenced using the indicated workflow.

**Supplemental Figure S4.**
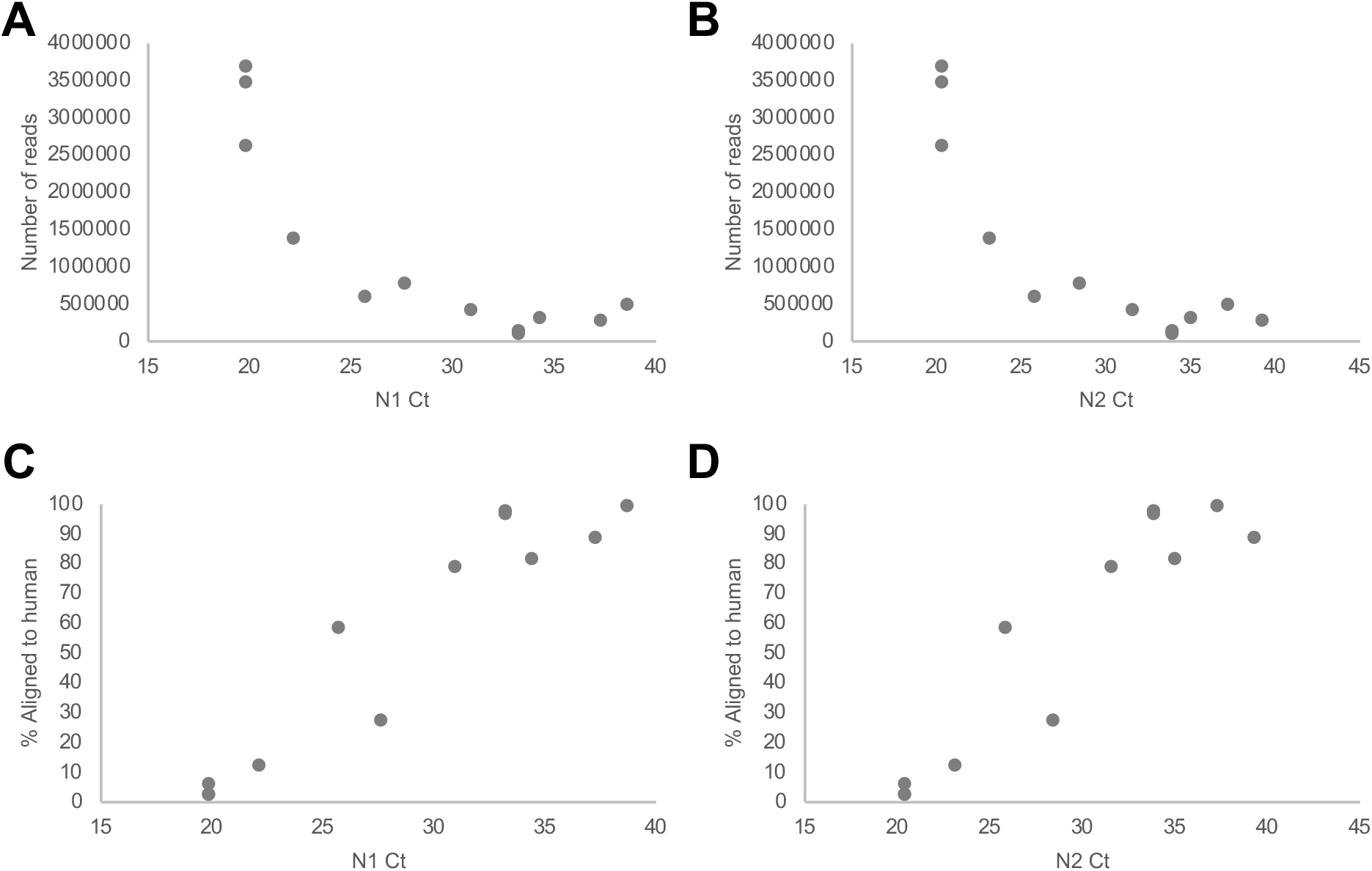
Performance metrics for Illumina DNA Flex Enrichment Protocol. Number of total reads generated per sample using the Illumina Nextera DNA Flex Enrichment workflow relative to: A) Sample N1 Ct value; B) Sample N2 Ct value. Percentage of reads aligned to a human reference genome using the Illumina Nextera DNA Flex Enrichment workflow relative to: C) Sample N1 Ct value; D) Sample N2 Ct value.

**Supplemental Figure S5.**
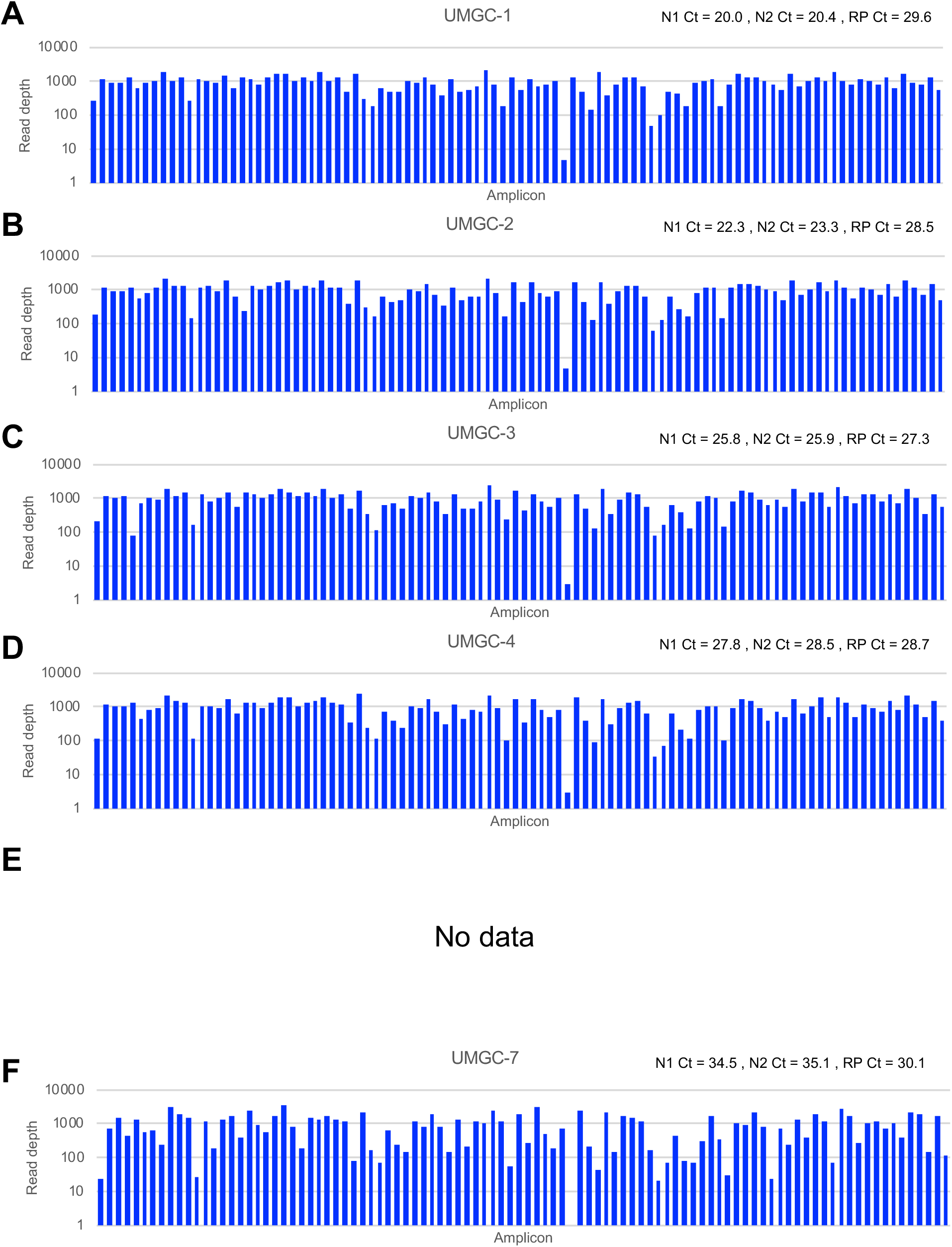
ARTIC v3 amplicon relative abundance. A-F) Observed read depth for each of the expected amplicons for the indicated sample amplified with the ARTIC v3 protocol with TruSeq library preparation at a subsampled read depth of 100,000 raw reads.

**Supplemental Figure S6.**
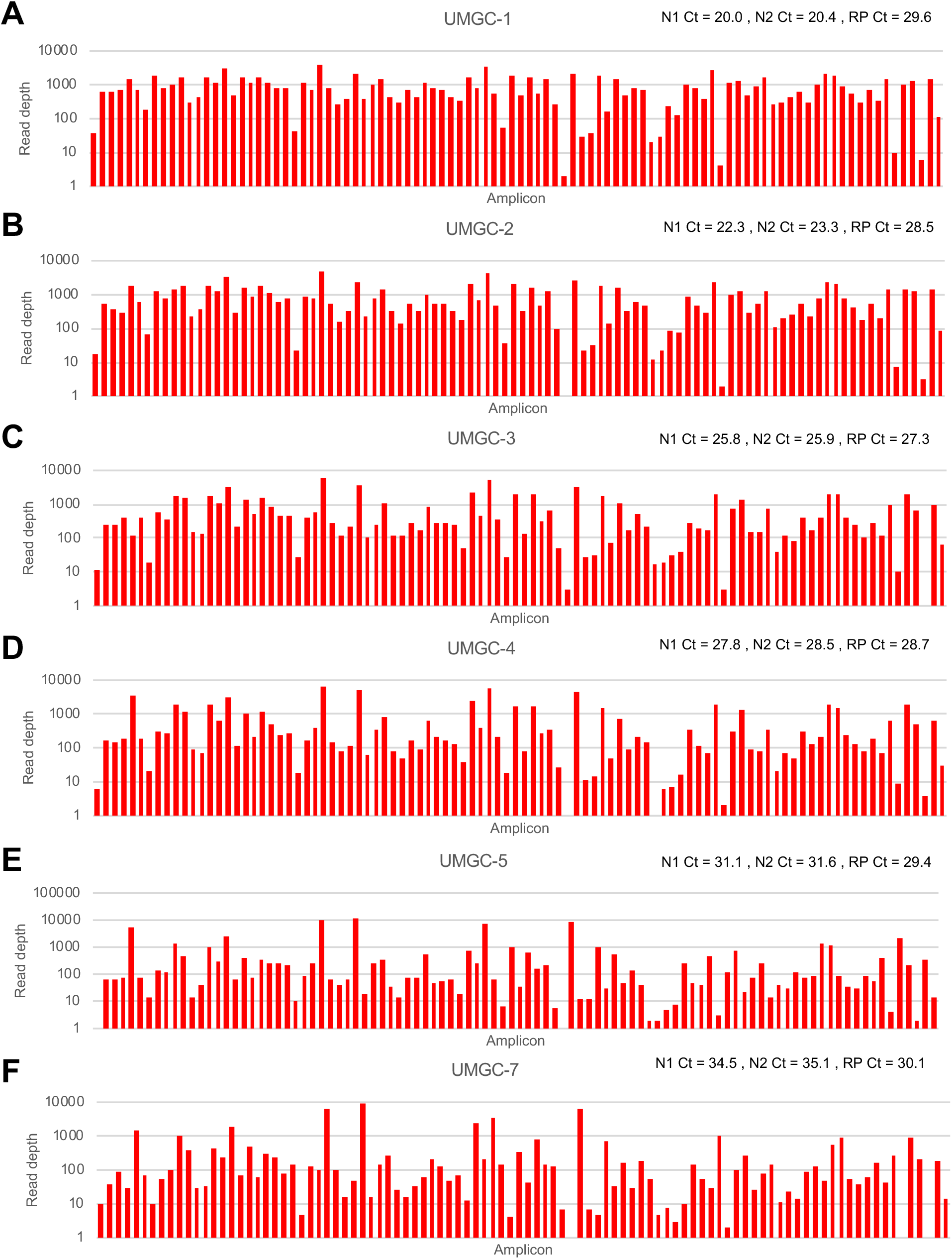
Tailed amplicon v1 amplicon relative abundance. A-F) Observed read depth for each of the expected amplicons for the indicated sample amplified with the tailed amplicon v1 protocol at a subsampled read depth of 100,000 raw reads.

**Supplemental Figure S7.**
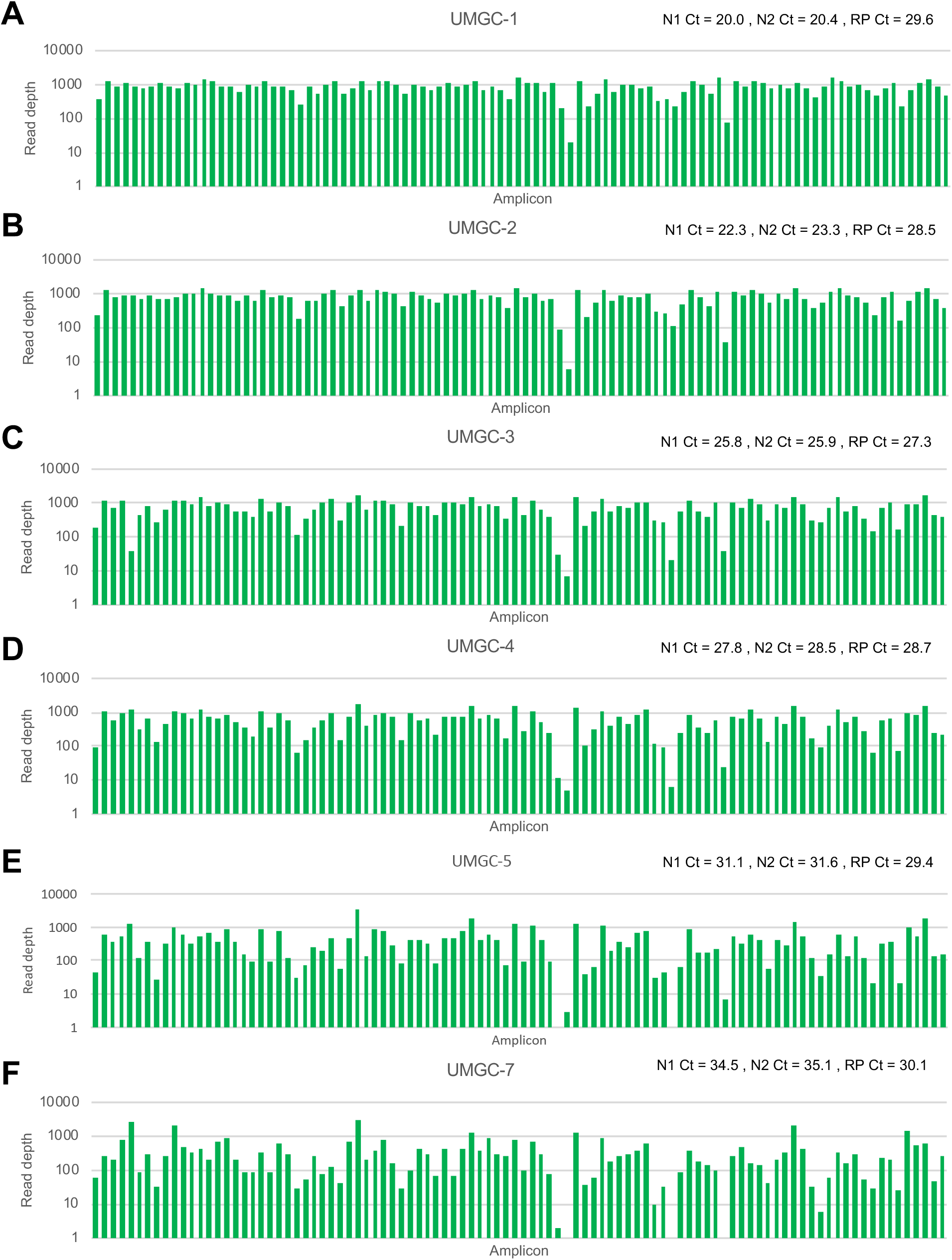
Tailed amplicon v2 amplicon relative abundance. A-F) Observed read depth for each of the expected amplicons for the indicated sample amplified with the tailed amplicon v2 protocol at a subsampled read depth of 100,000 raw reads.

**Supplemental Figure S8.**
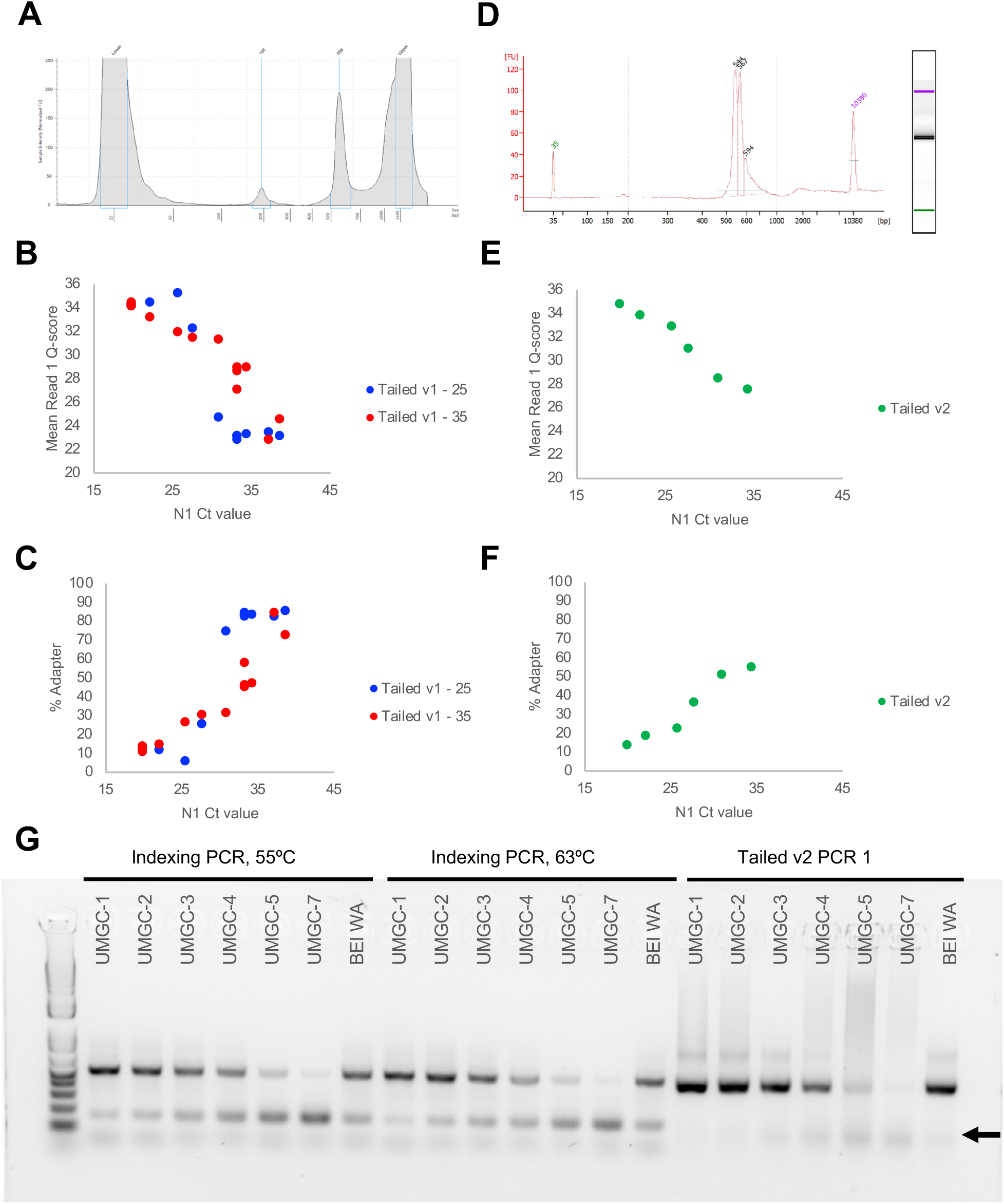
Primer dimer formation in tailed amplicon method. A) Agilent TapeStation trace for a library prepared from samples with N1 and N2 Ct values between ~20-40 using the tailed amplicon v1 (2 pool amplification) workflow. B) Mean read 1 quality score for samples prepared with the tailed amplicon v1 (2 pool amplification) workflow amplified for either 25 or 35 PCR cycles. C) Percentage of sequencing adapter observed for samples prepared with the tailed amplicon v1 (2 pool amplification) workflow amplified for either 25 or 35 PCR cycles. D) Agilent Bioanalyzer trace for a library prepared from samples with N1 and N2 Ct values between ~20-35 using the tailed amplicon v2 (4 pool amplification) workflow. E) Mean read 1 quality score for samples prepared with the tailed amplicon v2 (4 pool amplification) workflow. F) Percentage of sequencing adapter observed for samples prepared with the tailed amplicon v2 (4 pool amplification) workflow. G) 2% agarose gel showing the presence of primer dimers particularly in high N1/N2 Ct samples when indexed using different PCR cycling conditions. Arrow indicates primer dimers on gel.

**Supplemental Data File 1. Tailed amplicon v1 pool primer sequences** nCoV-2019_Nextera_UMGC_v1_pools.xlsx

**Supplemental Data File 2. Tailed amplicon v2 pool primer sequences** nCoV-2019_Nextera_UMGC_v2_pools.xlsx

